# Genetically encoded phase contrast agents for digital holographic microscopy

**DOI:** 10.1101/833830

**Authors:** Arash Farhadi, Manuel Bedrossian, Justin Lee, Gabrielle H. Ho, Mikhail G. Shapiro, Jay Nadeau

## Abstract

Quantitative phase imaging and digital holographic microscopy have shown great promise for visualizing the motion, structure and physiology of microorganisms and mammalian cells in three dimensions. However, these imaging techniques currently lack molecular contrast agents analogous to the fluorescent dyes and proteins that have revolutionized fluorescence microscopy. Here we introduce the first genetically encodable phase contrast agents based on gas vesicles, a unique class of air-filled protein nanostructures derived from buoyant microbes. The relatively low index of refraction of the air-filled core of gas vesicles results in optical phase advancement relative to aqueous media, making them a “positive” phase contrast agent easily distinguished from organelles, dyes, or microminerals. We demonstrate this capability by identifying and tracking the motion of gas vesicles and gas vesicle-expressing bacteria using digital holographic microscopy, and by imaging the uptake of engineered gas vesicles by mammalian cells. These results give phase imaging a biomolecular contrast agent, greatly expanding the capabilities of this powerful technology for three-dimensional biological imaging.

## INTRODUCTION

Precise acquisition of 4-dimensional data, comprising spatial coordinates and time, is important for studying many microscopic processes in biology. However, conventional optical microscopy suffers from a narrow depth of field due to its reliance on focusing lenses to encode spatial depth. Volumetric information is only obtained through step-wise acquisition of points or planes, and the resulting sequential acquisition typically necessitates low resolution in at least one of the 4 dimensions. Digital holographic microscopy (DHM), an inherently volumetric recording technique, provides an alternative for instantaneous sampling of thick volumes. Successful applications have been made to studies of microbial motility, chemotaxis, flow of ions through ion channels, and migration of cancer cells (*1–7*).

With DHM, optical interferometry is used to record a series of holograms at frame rates limited only by the camera. Off-line reconstruction yields plane-by-plane images of image intensity (brightfield) and phase. In DHM, as in other forms of quantitative phase imaging (QPI), phase contrast at any point *x, y* is related to the difference in indices of refraction between the medium (***n***_***m***_) and objects in the light path (***n***_***c***_) multiplied by the object height *h* (*8*):

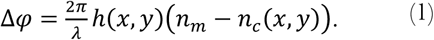

Because the typical values of *n*_*c*_ for the cytoplasm and organelles are approximately 1.38, which is very close to the *n*_*m*_ of 1.33 for water, it is challenging for DHM to visualize cells, particularly when they are small (*9*). This challenge could be overcome with suitable contrast agents or reporter genes, which would make cells more visible or highlight subcellular features and processes. In fluoresce microscopy, this function is provided by targeted small-molecule dyes and fluorescent proteins, which have revolutionized the utility of this technique in biological research (*10–13*). Unfortunately, these same molecules are not effective as phase contrast agents due to their small refractive index difference relative to H_2_O and its similarity to other intracellular materials (*14–18*). An ideal phase contrast agent would have a more dramatically different index of refraction, and preferably one that is lower than that of H_2_O to be categorically distinguishable from other cellular components.

Here we introduce genetically encodable contrast agents for phase imaging. These contrast agents are based on a unique class of hollow protein nanostructures called gas vesicles (GVs). GVs are all-protein nanostructures natively expressed in a number of waterborne microorganisms as a means to regulate their buoyancy (*19, 20*). GVs are air-filled compartments with dimensions on the order of 200 nm, enclosed by a 2 nm-thick protein shell (Fig. 1A), which is permeable to dissolved gas but prevents water from condensing into a liquid in the GV core. GVs have previously been described as contrast agents for ultrasound (*21*) and magnetic resonance imaging (*22–24*), but have not been applied to optical microscopy. Recently, the genes encoding GVs have been heterologously expressed in commensal bacteria (e.g. *Escherichia coli* Nissle 1917 and *Salmonella typhimurium*) (*23, 25*) and mammalian cells (*26*).

**Fig. 1.**
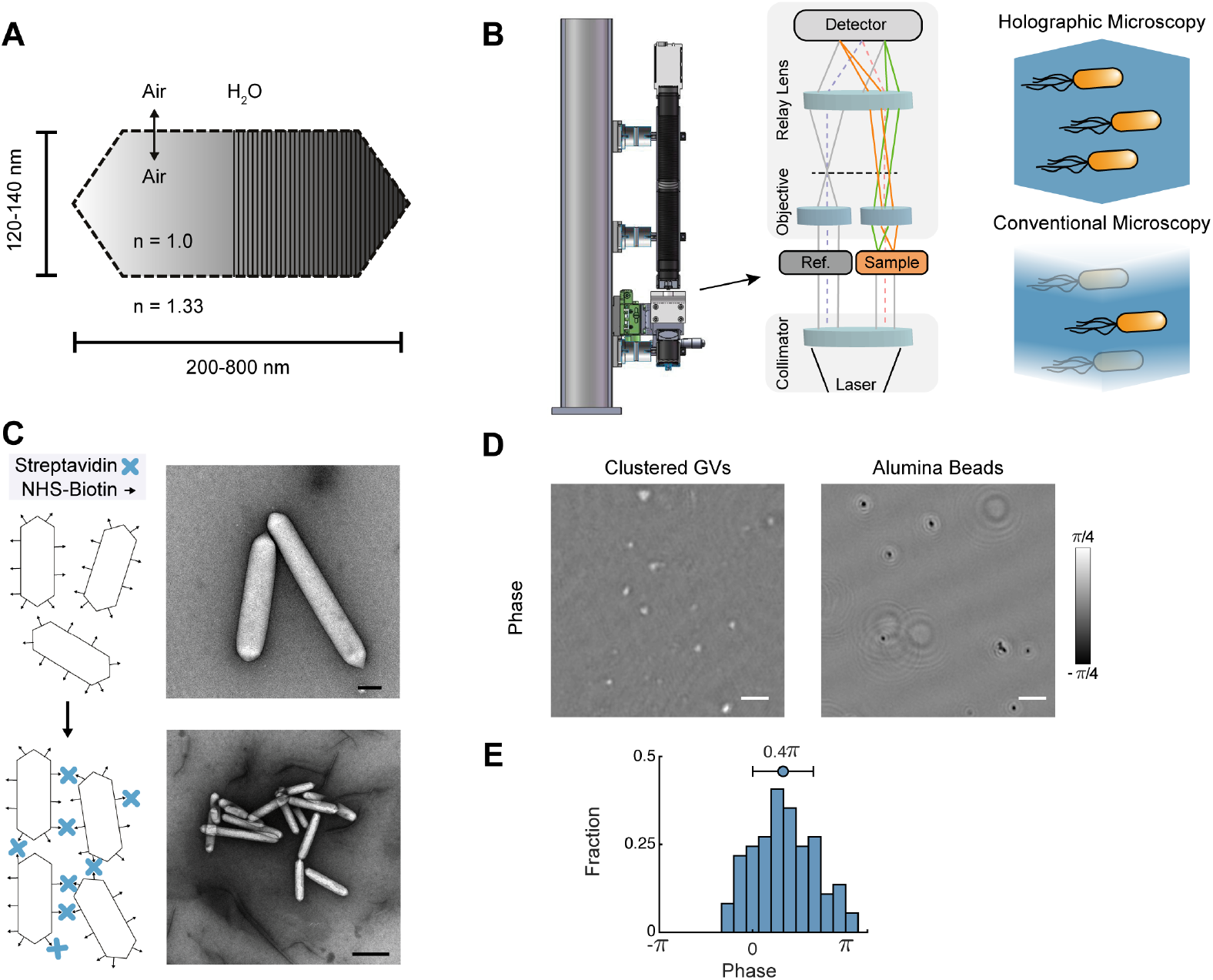
Gas vesicles as phase contrast agents. (A) Schematic of a single GV, indicating the index of refraction, n, inside the GV and in typical aqueous media. (B) Schematic of common path DHM system used in this work, and an illustration of the inherent volumetric imaging of DHM compared to conventional microscopy. (C) Biotinylated GVs purified from *Anabaena flos-aquae* can be clustered using streptavidin. Shown are a schematic and representative transmission electron micrograph of biotinylated GVs (top) and clustered GVs (bottom). Scale bars 100 nm and 500 nm, respectively. (D) Representative DHM images of clustered GVs and alumina beads. Scale bars represent 10 µm. (E) Histogram of phase change observed from clustered GVs with apparent diameters from 0.6-2.2 µm.

In this work, we test the hypothesis that the large difference in the index of refraction of GVs’ gaseous interior (n=1.0) relative to water and cytoplasm would produce strong positive phase contrast. We use a DHM system capable of providing sub-micron resolution to visualize the phase contrast produced by purified GVs, allowing 4-dimensional tracking of their motion. In addition, we show that *S. typhimurium* cells expressing GVs as a genetically encoded reporter produce a unique pattern of phase contrast reflecting the sub-cellular distribution of these nanostructures. Furthermore, we demonstrate the use of GVs as targeted molecular contrast sources in mammalian cells by visualizing the uptake of engineered GVs into a mammalian cell line. This work establishes GVs as a tool for molecular and genetically encodable phase contrast, greatly expanding the utility of DHM and related methods.

## RESULTS

### Gas vesicles produce positive phase contrast

To establish the ability of GVs to produce phase contrast, we imaged GVs purified from *Anabaena flos-aquae.* These nanostructures are approximately 120-140 nm wide and 200-800 nm long as measured by transmission electron microscopy (TEM) (Fig. 1A). Individually, purified GVs were too small to resolve on our DHM system, which uses numerical aperture (NA) of 0.3 aspheric lenses to achieve a spatial resolution of 0.8 µm (Fig. 1B). We therefore assembled GV clusters by biotinylation of the GVs followed by the addition of streptavidin. This yielded clusters with hydrodynamic diameters of 690±56 nm (Fig. 1C and **Supplemental Fig. 1**). Holograms were collected of GV clusters and of comparably sized alumina beads (1524±470 nm diameter as measured by dynamic light scattering) suspended in phosphate-buffered saline (PBS). We reconstructed these holograms into phase images as described in the supplementary text and **Supplemental Fig. 2**. The phase shifts from the GV clusters and alumina beads were opposite in sign, with the GV clusters exhibiting positive phase contrast, while the alumina beads exhibited negative phase contrast (Fig. 1D). The average phase shift produced by the GV clusters was 0.4 ± 0.32 π (Fig. 1E). GV phase contrast could be erased by irreversibly collapsing the GVs with hydrostatic pressure (*27*), providing a convenient internal control. After collapse, the positive phase contrast of GVs was eliminated (**Supplemental Fig. 3**).

Having established GVs as phase contrast agents, we sought to determine if DHM could be used to distinguish them and track their motion in a mixed particle population. First, we tracked the motion of both GV clusters and alumina beads in a 1 mm-deep well. The average speed of GV clusters rising along the depth axis due to their buoyancy was 0.43 ± 0.58 µm s^−1^ (95% C.I.) (Fig. 2A-C), while alumina beads sank with an average speed of –0.47±1.02 µm s^−1^ (95% C.I.) (Fig. 2D-F). When GV clusters and alumina beads were mixed together and tracked, the two particle types were readily distinguished (Fig. 2G-J), with the particles that were rising having positive phase contrast and the particles that were sinking having negative phase contrast (Fig. 2K). This demonstrated the ability of GVs to serve as a categorical contrast agent for 4-dimensional DHM.

**Fig. 2.**
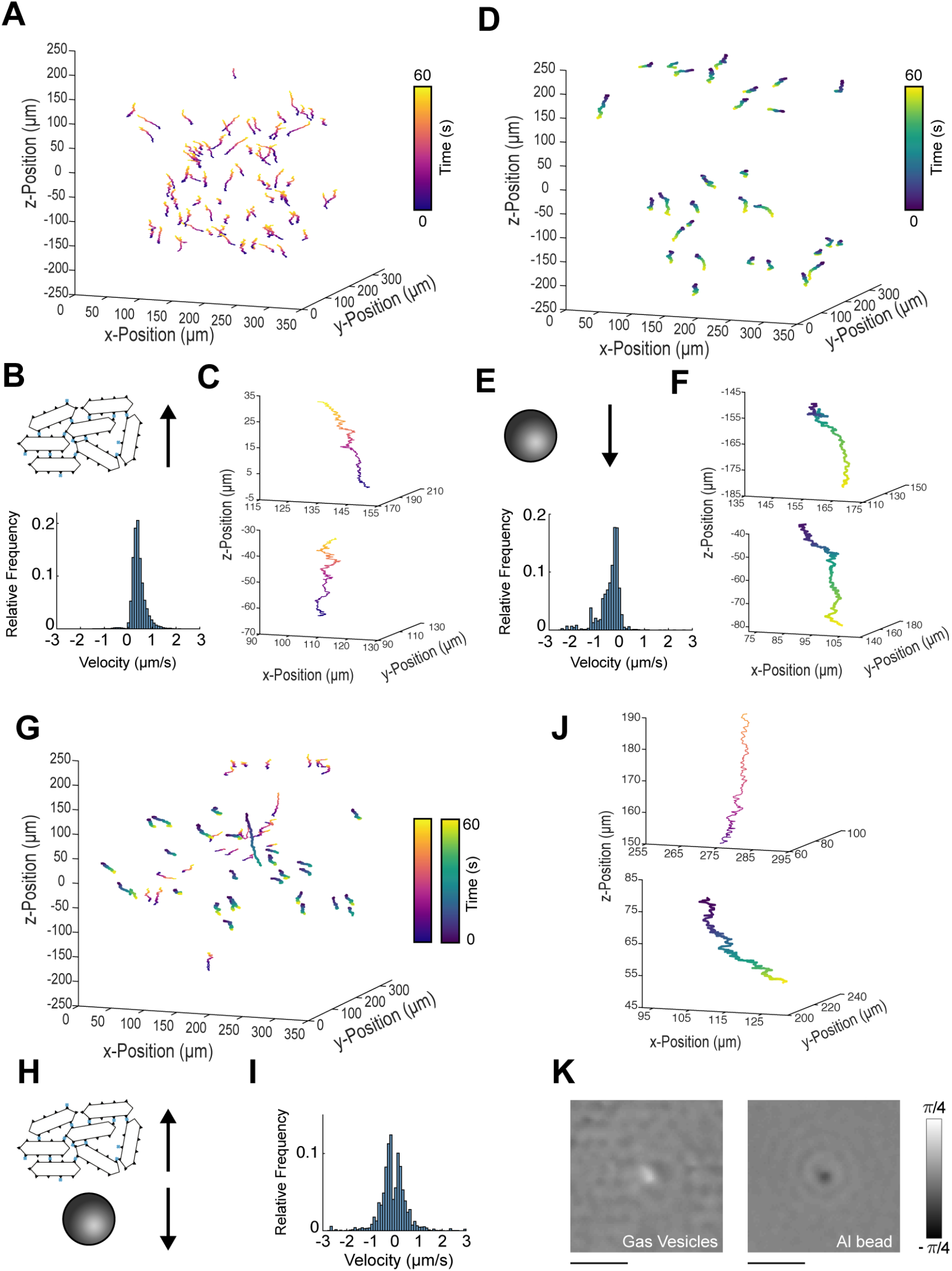
Volumetric tracking of particles in a mixed population suspension. (A) 3D trajectory plots of tracked GVs over 60 seconds. (B) Clustered GVs rise in solution (Top). Histogram of the z-component velocities of clustered GVs (Bottom). (C) Two example 3D tracks of clustered GVs. Each trajectory is color-coded with respect to time. (D) 3D trajectory plots of tracked alumina beads over 60 seconds. (E) Alumina beads sink over time (Top). Histogram of z-component velocities of alumina particles (Bottom). (F) Two example 3D tracks of alumina beads. Each trajectory is color-coded with respect to time. (G) 3D trajectory plots of the mixed population over 60 seconds. (H) Mixed population of clustered GVs and alumina beads. (I) Velocity histograms show two velocity populations, one for GVs and another for alumina beads. (J) Example trajectory of a rising and another of sinking particle chosen at random from G. (K) Phase images of the two particles from J. The buoyant particle has a positive phase contrast while the particle sinking has a negative phase contrast. Scale bars represent 5 µm.

### Engineered gas vesicles serve as targeted contrast agents

Among the advantages of GVs as phase contrast agents is their ability to be genetically engineered to modify their surface biochemical properties and enable molecular targeting (*27, 28*). To demonstrate that engineered, targeted GVs could be used for phase imaging in living cells, we used a fusion of the *A. flos-aquae* sequence to a polyarginine (R8) peptide. This peptide is a mimic of the human immunodeficiency virus 1 trans-activating (HIV-1 TAT) peptide and allows tagged proteins and particles to penetrate into mammalian cells (Fig. 3A) (*29*). Polyarginine-modified GVs were purified and added to Chinese hamster ovary (CHO) cells at 114 pM for 45 minutes, washed and imaged by a DHM system. This allowed for rapid 3-dimensional reconstruction of GV location on the surface of and within the cells (Fig. 3B & 3C). GVs were fluorescently labeled with Alexa Fluor 488-NHS ester dyes to independently confirm the location of GV labeling on the cells (Fig. 3D & 3E). These experiments demonstrate the ability of GVs to label living cells for 3-dimensinoal phase imaging.

**Fig. 3.**
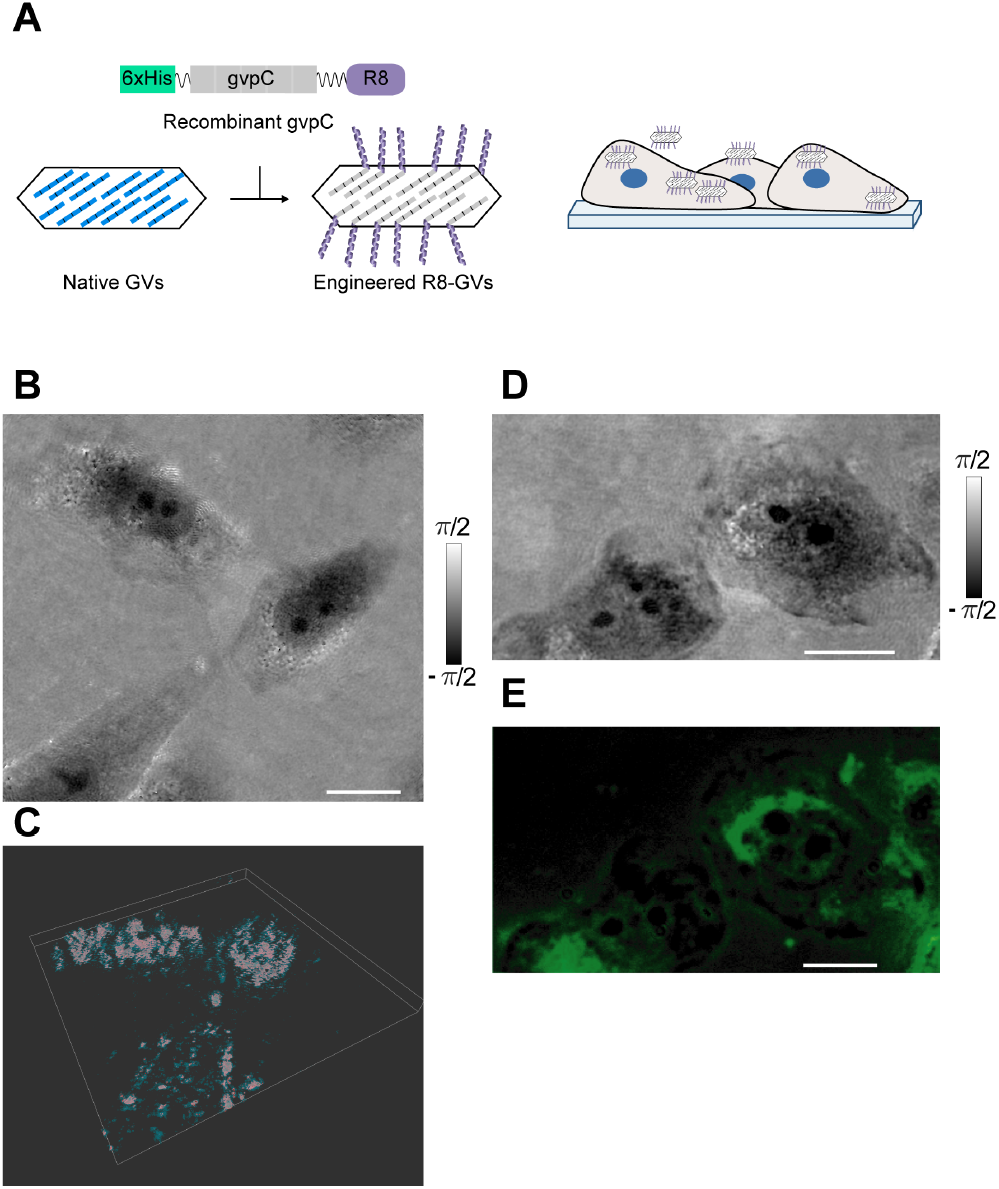
DHM imaging of mammalian cells labeled with engineered gas vesicles. (A) Diagram of engineered GVs genetically functionalized with R8 peptides for attachment to and internalization by mammalian cells. Illustration of mammalian cells labeled with R8-GVs. (B) DHM phase image of CHO cells labeled with R8-GVs. Scale bar represents 25 µm. (C) Pseudocolored 3D rendering of GVs decorating CHO cells. For details see supplemental text. (D) Phase image of CHO cells labeled with Alexa Fluor 488 conjugated R8-GVs under the high power DHM showing positive phase contrast in correspondence with (E) a fluorescence image of the same CHO cells. All scale bars represent 25 µm.

### Gas vesicles as genetically expressed contrast agents in Salmonella typhimurium

Following the characterization of GVs as targeted DHM contrast agents, we tested the ability of genetically encoded GVs to act as phase contrast reporter genes in living bacteria. A recently developed gene cluster encoding GVs, comprising a combination of genes from *Anabaena flos-aquae* and *Bacillus megaterium* (*25*), was used to heterologously express GVs in *Salmonella typhimurium* (Fig. 4A). Upon induction with *N*-(*β*-ketocaproyl)-*L*-homoserine lactone (AHL), *Salmonella* formed numerous intracellular GVs, which were visualized under TEM and seen to typically cluster into distinct subcellular regions (Fig. 4B). This pattern was expected to perturb the electromagnetic wavefront passing through a GV-expressing *Salmonella* cell so as to produce a distinct pattern in phase images, as shown in simulated holograms (Fig. 4C, Supplemental Text).

**Fig. 4.**
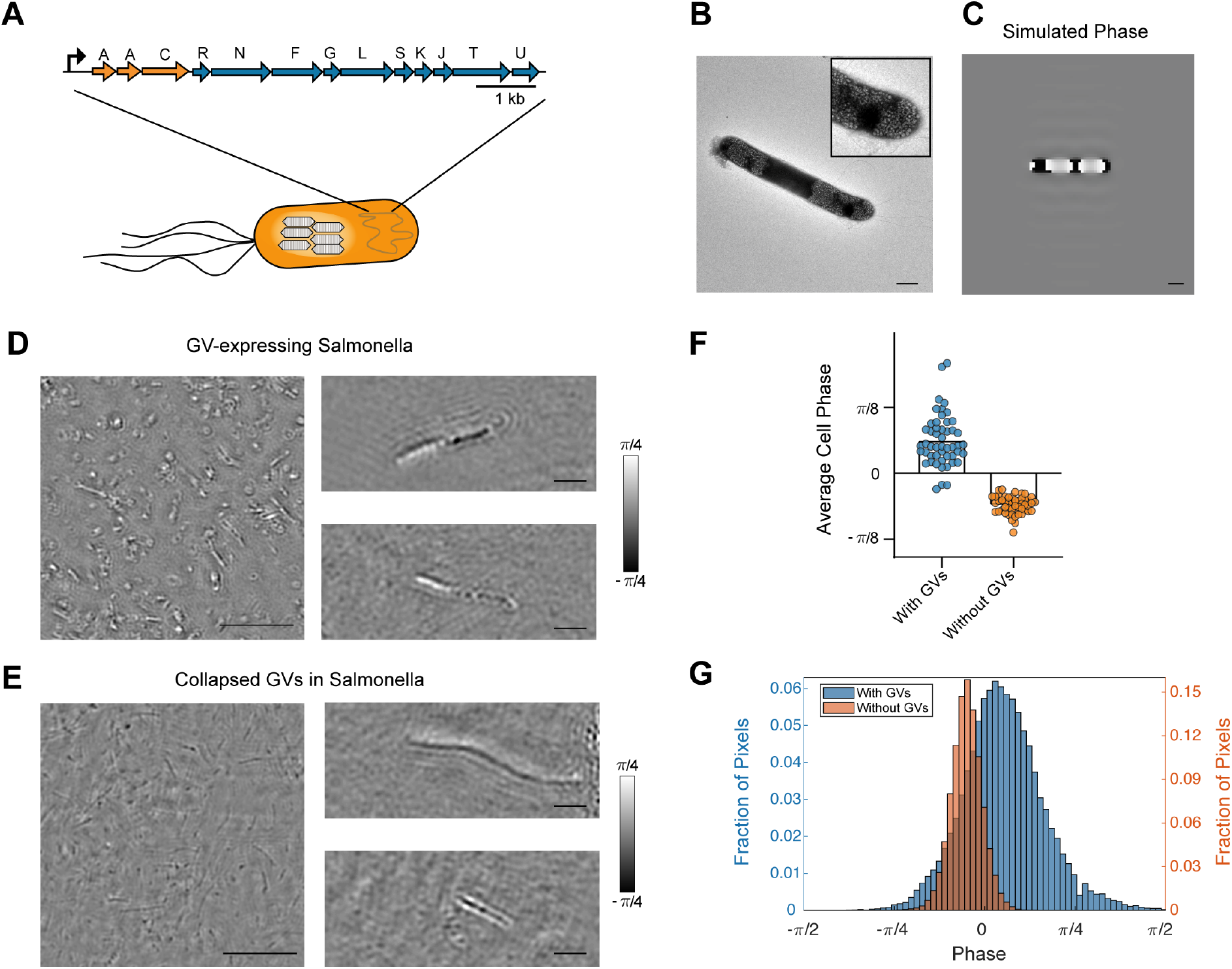
Gas vesicles as genetically encoded phase contrast agents in *Salmonella*. (A) Schematic of engineered GV gene cluster comprising genes from *Anabaena flos-aquae* (orange genes) and *Bacillus megaterium* (blue genes) that enable heterologous expression of GVs in *Salmonella typhimurium*. (B) Representative transmission electron micrograph of *S. typhimurium* expressing GVs. Inset is a 2x zoom in of a GV containing region of the salmonella cell. Scale bar represents 1 µm. (C) Numerical simulation of phase images of a GV-expressing *Salmonella* cell. Scale bar represents 1 µm. (D) Representative phase images of GV-expressing *S. typhimurium* cells under DHM. Two examples of zoomed in images are shown on the right. Scale bars for full field of view are 25 µm, and 5 µm for the zoomed in images. (E) Representative phase images of GV-expressing *S. typhimurium* cells after collapsing GVs using 1.2 MPa of hydrostatic pressure. Two examples of zoomed in images are shown on the right. Scale bars for full field of view are 25 µm, and 5 µm for the zoomed in images. (F) Quantified average phase contrast from *S. typhimurium* cells with intact GVs compared with collapsed GVs (n=50 cells). (G) Histogram of quantified phase contrast from all pixels in *S. typhimurium* cells with intact GVs (n=50 cells) compared with collapsed GVs (n=50 cells).

As expected, under DHM, GV-expressing *Salmonella* exhibited clear phase contrast that was different from control cells in which the GVs had been collapsed with hydrostatic pressure (*27*). Phase values of interior cellular regions of 50 GV-expressing *salmonella* cells were analyzed and found to exhibit an average phase value of 0.06 ± 0.05π (95% C.I., n=50 cells) (Fig. 4D & 4F). The subcellular regions of these cells which contained high GV concentrations exhibited even higher phase values of 0.42 ± 0.10π (95% C.I., n=110 sub-cellular regions). The phase values of interior cellular regions of 50 GV-expressing *Salmonella* cells after their GVs were hydrostatically collapsed were measured to be −0.06 ± 0.10π (95% C.I., n=50 cells) (Fig. 4E & 4F). The histogram of all pixels in *Salmonella* cells with and without GVs illustrates increased phase contrast due to GV-expression (Fig. 4G). GV-expression increased the average signal to noise ratio of *Salmonella* cells in phase images by more than three-fold, making the cells easier to detect in solution.

## DISCUSSION

Our results demonstrate that GVs serve as molecular and genetically encodable contrast agents for phase imaging due to the large difference in their index of refraction compared with aqueous media and organelles, resulting in positive phase contrast. The opposite sign of their refractive index difference from water compared to most other biological structures is especially beneficial for studying small samples such as microorganisms or subcellular features. The rapid volumetric image acquisition of DHM makes it possible to identify and dynamically track GVs in a mixed solution containing other particles. In addition, the modular protein make-up of GVs enables protein engineering to confer novel targeting properties for cellular imaging. This may facilitate future studies of cell-nanoparticle interactions and other dynamic cellular processes. Furthermore, by introducing GV gene clusters into engineered cells, we can obtain genetically encoded phase contrast, allowing cells activated to express these reporter genes to be distinguished from cells without reporter expression. These studies result in a QPI toolkit that will permit specific labeling in a large number of biological scenarios.

While the results presented in this study provide the key scientific evidence supporting the ability of GVs to serve as genetically encoded reporters for phase imaging, future work is needed to apply these agents to specific biological problems. To enable such applications, there exists significant scope for improvement and optimization. In using GVs as a contrast agent for arbitrary samples, phase wrapping must be considered. In a typical QPI system, phase measurements are constrained to a modulo-2π (e.g. -π to π). As a result, samples that introduce a phase shift of ***ϕ* = 2*π* + *m***, will only be seen to show a phase shift of ***m***. Targeted efforts in the development of robust phase unwrapping algorithms will aid in the use of GVs in arbitrarily thick samples, eliminating the loss of contrast due to aliasing (*30–35*). Given the ability of GVs to be collapsed with acoustic pressure, additional contrast specificity could be obtained by integrating the ability to apply ultrasound waves to DHM samples *in situ* and monitoring the resulting change in phase (*23, 25, 27*). In addition, the engineering of GV variants that collapse under different applied pressures may enable multiplexed phase contrast imaging. Furthermore, while genetic encoding facilitates the use of GVs to study genetically tractable organisms, there is also substantial interest in using DHM for field studies, taking advantage of the microscopes’ robust solid-state design (*36*). In such studies, the relevant microorganisms may be genetically intractable, requiring the development of targeting moieties to bind GVs to such species. Such labeling would additionally facilitate the application of machine learning algorithms to the detection of microorganisms in 4-dimensional data, where low image contrast is currently a challenge (*37*). We anticipate that the development of dedicated molecular and genetically encodable contrast agents will usher in a new phase in holographic microscopy.

## MATERIALS AND METHODS

### Digital holographic microscopy

Two instruments were used: a “high power” instrument and a “low power” instrument. The design of the “high power” microscope was a modified Mach-Zehnder as described previously (*38*), containing identical objective lenses in the object and reference beams. The objective lenses used were Mitotoyo 100x, NA 0.7 dry long working distance objectives, infinity-corrected to an achromatic field lens (200 mm focal length), which was used to form the image on a digital CCD camera (Baumer TXG50-P). The effective magnification of this instrument was 78x. The diffraction-limited lateral resolution was roughly 0.3 µm with a 405 nm illumination source. Illumination was through a single-mode fiber coupled diode laser that was collimated before the first beamsplitter.

The design of the “low power” microscope was a common path Mach-Zehnder as described previously (*36, 39*). The objectives were simple aspheric lenses (Asphericon) with NA = 0.3. The effective magnification was 19.6x with a diffraction limited lateral resolution of 0.8 µm. The wavelength used in this work for both DHM instruments was 405 nm, supplied by a diode laser (Thorlabs S1FC405).

DHM images of GV-labeled mammalian cells were acquired using the “high power” microscope and all other DHM data were collected using the “low power” microscope.

### Gas vesicle expression, purification and clustering

*Anabaena flos-aquae* (CCAP strain 1403/13F) was cultured for ~2 weeks in Gorham’s media supplemented with BG-11 solution (Sigma) and 10 mM NaHCO_3_ at 25ºC, 100 rpm under 1% CO_2_ and a 14 hours light/10 hours dark cycle (*28*). At confluency, the buoyant cell fraction was isolated by floating to the top of a separating funnel over a 48h period, after which the subnatant was discarded. The collected cells were then lysed using 500 mM sorbitol and 10% Solulyse solution (Genlantis). GVs were purified through repeated rounds of isolating the buoyant fraction through centrifugation and resuspension in PBS.

GV clusters were prepared by first biotinylating purified GVs with 10^5^ molar excess of EZ-Link Sulfo-NHS-LC-biotin (Thermo Scientific) in PBS for 4 hours. Afterwards, the sample underwent two rounds of overnight dialysis in PBS. The biotinylated GVs were then clustered by incubation with streptavidin (Geno Technology) for 30 minutes at room temperature at a streptavidin to GV molecular ratio of 100:1.

### Dynamic light scattering

The hydrodynamic size of the GVs, GV clusters and alumina beads was measured in 50 µL samples at OD_500_ = 0.2 using a Zeta-PALS analyzer (Brookhaven Instruments). Samples were mixed thoroughly and measured five times for each reported hydrodynamic diameter.

### Engineered gas vesicles for cell labeling

Genetically engineered GVs were prepared using a previously described protocol (*27*). In brief, the GvpC DNA sequence from *Anabaena flos-aquae* was codon-optimized for *E. coli* expression and cloned into a pET28a(+) plasmid (Novagen) with an N-terminal hexahistidine-tag and C-terminal GSGRRRRRRRR sequence. Plasmids were transformed into BL21(DE3) cells (Invitrogen), which were induced to express the recombinant GvpC for 6 hours at 30°C. GvpC contained in inclusion bodies was purified by lysing the cells using 10% Solulyse (Genlantis) supplemented with DNaseI (10 μg/mL) and lysozyme (400 µg/mL) at room temperature. Inclusion bodies were recovered by centrifugation at 27,000g for 15 min. The inclusion body pellets were resuspended in 20 mM Tris-HCl buffer with 500 mM sodium chloride and 6 M urea (pH=8.0) and incubated with Ni-NTA resin (Qiagen, Valencia, CA) for 2 hours at 4°C. After washing, proteins were eluted using 250 mM imidazole.

Purified GVs were treated with 6 M urea and 20 mM Tris-HCl (pH=8.0) to remove their wild-type GvpC. The stripped GVs were isolated with two rounds of centrifugally assisted buoyancy purification in urea. Purified polyarginine modified-GvpC was then added according to the formula: 2 x optical density x 198 nM x liters GVs = nmol of recombinant GvpC and dialyzed in PBS for 8 hours. 10^5^ molar excess of Alexa Fluor 488 NHS (Thermo Fisher) was then added to the GVs and incubated at room temperature for 4 hours under gentle rotation, before being quenched with 20 mM Tris-HCl (pH = 8.0) and dialyzed in PBS to remove excess dye.

### Cell culture and gas vesicle labeling

Chinese hamster ovary cells (CHO-K1; ATCC) were cultured in DMEM (Corning) with 10% FBS (Thermo Fisher) and penicillin/streptomycin (Corning). Coverslips (18×18 mm) were sterilized with 70% ethanol, washed twice in PBS and placed in 6-well plates. Fibronectin (Sigma) was diluted 1:20 in PBS and 200 µL were added to each well over the entire surface of the coverslip and incubated for 1 hour at room temperature. Excess solution was aspirated, and CHO-K1 cells were seeded on the coverslips and grown to ~75% confluency.

For GV labeling, the surface of a 6-well plate was covered with paraffin and UV sterilized. Then, 300 µL of 37°C DMEM media and 300 µL of 114 pM (36.6 µg/mL or OD_500_ = 1) of R8-GVs was added to the bottom of the well and mixed. The cells cultured on coverslips were inverted onto the DMEM and GV mixture, so the cells were facing the bottom of the plate. The coverslips and GVs were incubated at 37°C. Following incubation, the cells were washed three times with 200 µL of PBS and fixed with 1 mL of formaldehyde for 30 minutes. The coverslips were mounted using Diamond Antifade mountant (Thermo Fisher) and sealed using clear nail polish.

### Fluorescence imaging

Fluorescence images were taken on an Olympus IX-71 inverted microscope using Hg lamp illumination through a 1.4 NA oil immersion objective and using the enhanced green fluorescent protein filter set (Chroma). To register fluorescence images with phase images, a 1-µm tip glass pipette was secured to the specimen and cells were imaged in the vicinity of the tip across the two instruments.

### Gas vesicle expression in Salmonella

GV expression in *Salmonella typhimurium* (strain ELH1301) cells was performed as described previously (*25*). Briefly, the hybrid GV gene cluster, under the control of the *luxI* promoter (addgene 106475), was transformed into *S. typhimurium* cells. Monoclonal cells from an individual plated colony were cryostocked. Cells containing the GV genes were grown in 5 mL of 2xYT media with 50 µg/mL kanamycin for 16 hours at 37ºC, 250 rpm. Cultures in 50 mL 2xYT media with 50 µg/mL kanamycin were then inoculated with 500 µL of the starter culture and grown on the shaker at 37ºC until OD_600_ = 0.4 to 0.6. These cultures were induced with 3 nM N-(β-ketocaproyl)-L-homoserine lactone (AHL) and then grown for 22 hours at 30ºC, 250 rpm. Cells were then harvested by centrifugation at 300g at 30ºC for 2 hours. The buoyant cell fraction was transferred into clean tubes. To collapse the GVs inside cells, GV-expressing salmonella were placed a quartz cuvette (Hellma Analytics) connected to a N_2_ cylinder through a pressure controller (Alicat Scientific) set to 1.2 MPa.

### TEM sample preparation and imaging

Electron microscopy was performed at the Beckman Institute Resource Center for Transmission Electron Microscopy at Caltech. Purified GVs were diluted to OD_500_ = 0.2 in 10 mM HEPES buffer and *Salmonella* cells were diluted to OD_600_ ~ 0.2 in 10 mM HEPES buffer or PBS. Samples were then spotted on Formvar/Carbon 200 mesh grids (Ted Pella), which were rendered hydrophilic by glow discharging (Emitek K100X). Purified GV samples were stained with 2% uranyl acetate, while cells were imaged unstained. Image acquisition was performed using a Tecnai T12 Lab6 120 kV transmission electron microscope equipped with a Gatan Ultrascan 2k x 2k CCD camera.

### Simulations

Holograms were simulated with MATLAB (R2017b) using a custom hologram simulation routine. First, a two-dimensional projection image was created using a series of Radon Transforms, modeling a typical GV-expressing *Salmonella* cell as a cylinder, with a diameter of 1 µm and a length of 5 µm, with bands of lower refractive index corresponding to areas dense in GVs as seen in TEM images. This projection is then downsampled, via bicubic interpolation, in order to accommodate and emulate the diffraction limited resolution of the low-power DHM instrument. Within this projection, GVs were simulated using index of refraction values of 1.00. Other intracellular areas were simulated using an index of refraction of 1.37. The index of refraction used to simulate the cell’s surrounding medium is 1.33.

The wavefront perturbations as a result of a collimated plane wave passing through the simulated cell is then propagated using the angular spectrum method (*40*). The resulting diffracted wavefront is numerically propagated and recombined with a reference (undisturbed) plane wave in order to simulate an off-axis hologram. Code for the simulator is provided in the Supplemental Material. The simulated holograms were reconstructed into phase images using the commercially available software KOALA (LynceeTec). No image noise was added to the simulation besides quantization noise when the holograms were saved as unsigned 8-bit image files whereas in reality there are numerous sources of noise including, but not limited to, photon ‘shot’ noise, temporal and spatial noise due to changes in the coherence of the illumination laser, as well as various sources of noise introduced by the digital CCD used to record the holograms(*9*).

### Tracking

Tracking of GVs and alumina beads was performed using the Manual Tracking plug-in in the open source image analysis tool FIJI (*41*).

### Phase quantification of Salmonella cells

Data recorded using the DHM system was reconstructed into phase images using the commercially available software Koala (LynceeTec). Raw 8-bit phase images were reconstructed with quantitative phase bounds of –π to π corresponding to pixel values of 0 and 255, respectively (described in supplementary text). After reconstruction cell boundaries were identified with the freehand selection tool of the open source image analysis software FIJI by team members blinded to the identity of the sample. The cell boundaries were used to isolate the interior pixel values within the cell by creating a binary mask about the cell boundary. These interior pixel values were converted from their 8-bit values to quantitative phase values and analyzed using MATLAB (2017b).

## ACKNOWLEDGEMENTS

The authors would like to thank Raymond Bourdeau for assistance with initial experiments, Kurt Liewer for the hologram simulation code and Dina Malounda for providing some of the samples. Arash Farhadi was supported by the NSERC graduate fellowship. Research in the Shapiro lab was supported by the National Institutes of Health (R01EB018975), the Packard Fellowship in Science and Engineering, the Pew Scholarship in the Biomedical Sciences and the Heritage Medical Research Institute. The Nadeau lab was supported by the Gordon and Betty Moore Foundation as well as the JPL-NEXT program.

## AUTHOR CONTRIBUTIONS

AF, MB, MGS, and JN conceived and planned the research. AF, JL and GH prepared the biological samples. MB and JN conducted the DHM data collection and reconstruction with help from AF. MB carried out the generation and analysis of the salmonella hologram simulations as well as the analysis particle and salmonella image reconstruction and data analysis. MB performed the processing and rendering of the 3D pseudo-colored phase image of CHO cells. AF, MB, MGS, JL, GH and JN analyzed the results. AF, MB, MGS and JN wrote the manuscript with input from all authors. MGS and JN supervised the research.

