## Supplementary Text and Materials for "Genetically encoded phase contrast agents for digital holographic microscopy"

#### Supplementary Figures and Tables

### Image Processing and Handling

In the analysis of off-axis holograms, image reconstruction and post-processing steps are necessary in order to interpret the electric field intensity recorded in a hologram into a useful three-dimensional data set. Furthermore, post-processing methods are used to de-noise the resulting reconstructed images.

This supplemental document outlines the methods and work flows associated with the processing and handling of off-axis holographic images. This includes the reconstruction process from hologram to phase images, and all de-noising steps used to reach the final images that are presented in the main document.

##### Image Reconstruction

An off-axis hologram is recorded as a single image that contains interference patterns (or fringes) that contain the 3D information of the sample being imaged. The spatial frequency of the fringes act as the carrier frequency of this information. To reconstruct this information into a useable form, the high spatial frequency information in the hologram must be isolated. **Supplementary Figure 1A** shows an example hologram, its Fourier Transform plotted on a logarithmic scale (note the high frequency 'lobes' which contain the 3D information of the sample (**Supplementary Figure 1B**), the same Fourier transform that has been multiplied by a binary mask to isolate one of the high frequency lobes (**Supplementary Figure 1C**), and that isolated lobe that has been shifted to the center of the Fourier Transform (**Supplementary Figure 1D**).

In the formation of an off-axis hologram, two collimated beams of light are recombined at the digital detector at an angle (off-axis of each other). This off-axis recombination causes interference between the two beams. The spatial frequency of these fringes is proportional to the wavelength of light used as well as the recombination angle. If the two beams of light are named the 'specimen' and 'reference' beam where the 'specimen' beam interacts with the sample being imaged, and the 'reference' beam remains undisturbed, then the resulting hologram can be represented mathematically as the superposition of the two beams at the detector (Equation 1).

$$S(x,y) = A_S \exp i(\phi_S(x,y) - \omega t), \quad \text{1a}$$
$$R(x,y) = A_R \exp i(\phi_R(x,y) - \omega t), \quad \text{1b}$$
$$\Psi(x,y) = S(x,y) + R(x,y) \quad \text{1c}$$
$$h = \int \Psi\Psi^* \mathrm{d}t = (I_S + I_R) + SR^* + S^*R, \quad \text{1d}$$

where $A_S$ and $A_R$ are the amplitudes of the Specimen and Reference beam, $\phi_S$ and $\phi_R$ are the phase differences between the specimen and reference beams, $\Psi$ is the resulting wave from the superposition of the specimen and reference beams, and $h$ is the hologram.

In Supplementary Figure 1B there are three discrete lobes present. The center most lobe corresponds to the summation of intensities of the specimen and reference beams $(I_S + I_R)$ from equation 1d, while the two higher spatial frequency lobes correspond to $SR^*$ and $S^*R$ from Equation 1d. These two lobes are complex conjugates of each other, but for the purposes of this work, the lobe corresponding to $SR^*$ is the chosen lobe to be isolated in Supplementary Figure 1C & 1D.

With the 3D information encoded in the hologram isolated, it is then convolved with the discrete solution to the Fresnel Diffraction Integral ($G$) as described by Schnars, et al.(*1*) The reconstructed complex wavefront $\Gamma$ is

$$\Gamma(x,y,z,t) = \mathfrak{F}^{-1}[\mathfrak{F}\big(h(x,y,t)\big) * G(z)], \quad \text{2a}$$
$$G(z) = \exp\left\{\frac{-2\pi i z}{\lambda}\sqrt{1 - \frac{\lambda^2\left(n + \frac{N^2\Delta x^2}{2z\lambda}\right)^2}{N^2\Delta x^2} - \frac{\lambda^2\left(m + \frac{N^2\Delta y^2}{2z\lambda}\right)^2}{N^2\Delta y^2}}\right\} \quad \text{2b}$$
$$A(x,y,z,t) = |\Gamma(x,y,z,t)| \quad \text{2c}$$
$$\phi(x,y,z,t) = \arctan\left(\frac{\Im(\Gamma(\mathrm{x,y,z,t}))}{\Re(\Gamma(\mathrm{x,y,z,t}))}\right), \quad \text{2d}$$

where, $\mathfrak{F}$ is the Fourier Transform operator, the amplitude image ($A$) is calculated as the magnitude of the complex wavefront $\Gamma$, and the quantitative phase image ($\phi$) is defined as the inverse tangent of the imaginary parts of $\Gamma$ divided by the real parts of $\Gamma$. The function $G$ is a pure phase object that describes the propagation of an electric field through the focal plane ($z$) and can be modulated to calculate the reconstructed wavefront throughout an entire volume. For a more in-depth derivation of this function, see Schnars, et al.

**Post-Processing**

With the reconstructed amplitude $A(x,y,z,t)$, and phase images $\phi(x,y,z,t)$, post-processing is necessary to eliminate noise from various sources including, but not limited to, photon/shot noise, speckle noise, digitization noise, as well as detector noise. In addition, low spatial frequency artifacts are also common in phase reconstructions due to tilt in the sample chamber relative to the optical axis as well as objective lens curvature.

To remove as much noise from the reconstructed images as possible, temporally averaged images are calculated and subtracted from each phase image, as well as conducting spatial frequency band-pass filtering.

In the subtraction of temporally averaged phase (referred to as ‘mean subtraction’), the mean of each pixel is calculated through time. This mean image is then subtracted from the image used to calculate that mean. This effectively removes any stationary artifacts from the image highlighting any transient particle in the image.

After this mean subtraction, band-pass filtering is done to remove any low and high spatial frequency noise from the images. Low spatial frequency noise can be caused by tilt in the sample chamber relative to the optical axis, as well as by the curvature of the lenses used in the DHM instrument. Because the DHM is an instrument that is capable of achieving diffraction limited resolution, there is a physical limit of the spatial frequencies that can be recorded. This presents a clear upper limit to the spatial frequencies that carry useful information in the image. Any higher frequency artifacts in the image are by definition pure noise and are filtered out.

Because the phase images obtained using off-axis DHM are of a quantitative nature, this is the extent of the post-processing performed in order to preserve the quantitative information contained in the image. Other methods are more useful in enhancing contrast and suppressing noise but were not performed in this work.

**Physical Properties**

| **Property** | **GV Cluster (air)** | **Alumina** | **Water** |
|---|---|---|---|
| **Density ($kg/m^3$)** | 1.2 | 2700 | 1000 |
| **Viscosity ($N\ s/m^2$)** | - | - | $8.9\times10^{-4}$ |

#### Hologram Simulator

The optical theory used in the MATLAB code discussed in this document are well reviewed by Ulf Schnars, et al.(*1*). The main function (‘OAhologramSimulator.m’) expects as an input, two variables. The first is called ‘waveFront’. This is a complex matrix of the size of the final image. This matrix describes the normalized amplitude and phase of the electric field of the sample we wish to simulate. For the purposes of this manuscript, the variable ‘waveFront’ is provided in the file ‘waveFront.mat’. The second input variable is ‘desiredFileName’, which is the desired file path and name of the output. The output is the final 8-bit TIFF hologram.

The variable ‘waveFront’ is generated by creating two three dimensional matrices of the electric field attenuation and the index of refraction of light that passes through a cylindrical simulated ‘salmonella cell’ with the morphology as shown in the main manuscript.

With the two 3D matrices, a series of radon transforms are used to project a collimated beam of light through the object. These projections are then used to calculate the resultant wave front as a result of the projected electric field attenuation and phase delay (introduced by the index of refraction matrix). The hologram simulation code can be separated and described in eight sections.

**1 DHM Parameters**

This section establishes the optical performance parameters of the DHM instrument that is to be simulated. In this implementation, the off-axis holograms that are being simulated are of the 'low-power' instrument and have the appropriate performance parameters for that instrument.

Note that the numerical aperture of the objective lenses is not necessary because the geometric properties of the objective lens is inputted as 'fl' and 'DiaLens' (lens focal length and lens diameter, respectively).

**2 Create the undisturbed reference beam**

This section creates the reference wave. The reference wave by definition is undisturbed and un-attenuated and thus the reference wave is a simple matrix of ones.

**3 Propagate target and ref to lens object focal plane**

This section takes the input variable 'waveFront' and the newly created reference wave front 'U2ref' and propagates the two waves to the focal plane of the objective lens. The propagation of the two waves is conducted using the Angular Spectrum Method (ASM) as described in Schnars, et al. The propagation is done by calling an external function 'ang_spec_prop.m'. The output of this section are the variables 'U3' and 'U3ref', corresponding to the propagated wave front of the sample (U3) and reference (U3ref).

**4 Propagate to the lens**

This section takes the output from the previous section and propagates the two wave fronts to the objective lens. This section uses the same external function as the previous section. The outputs are 'U4' and 'U4ref'.

**5 Simulate the phase delay introduced by the lens**

This section models an ideal objective lens that introduces a phase delay into light as it travels through the lens. This simulates an ideal lens because it introduces no wave front aberrations or electric field attenuation.

In addition to simulating the phase delay caused by the objective lens, it applies this phase delay to 'U4' and U4ref'. The output of this section is 'Alens' and 'Alensref'.

**6 Propagate to focal plane**

This section takes the output of the previous section and propagates the two wave fronts to the back focal plane of the objective lens. The outputs of this section are the variables 'U5' and 'U5ref'.

**7 Combine wavefront with reference wave**

This section acts as the relay lens of the 'low-power' instrument by recombining the object wave front (U5) with the reference wave front (U5ref) as well as introducing a tilt angle between them so that they create interference patterns at the 'detector'. The output of this section becomes a single complex wave front 'Uccd'.

**8 Generate and save the hologram**

Because optical detectors such as CCD's only image electric field intensities, the intensity of 'Uccd' is calculated and saved as an 8-bit TIFF image with the file path and location inputted at the very beginning of this routine.

#### 3D Pseudo-Colored Rendering

The 3D rendering was generated using the commercially available software ARIVIS Vision4D. This software allows the import of a multidimensional image that is to be visualized in a variety of ways. The 2D images that comprise the 3D image stack used in ARIVIS were first processed using the methods described in the Image Processing & Handling section of this document. These processed images were then thresholded using a user defined threshold. Next the magnitude of the image gradient was calculated and stored as a separate stack of 2D images. The two 3D image stacks were then imported to ARIVIS Vision4D. The pixel values were plotted using a pseudo-colored lookup table. The opacity of each pixel was plotted as a weighted function of the magnitude of the image gradient at that pixel. This was done to highlight areas of large changes in phase signal within the cell.

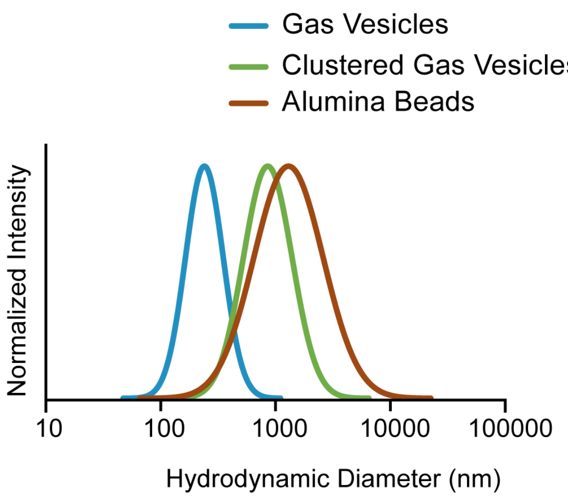


**Supplementary Figure 1 -** Representative dynamic light scattering of the hydrodynamic diameter of pristine gas vesicles, clustered gas vesicles and alumina beads.

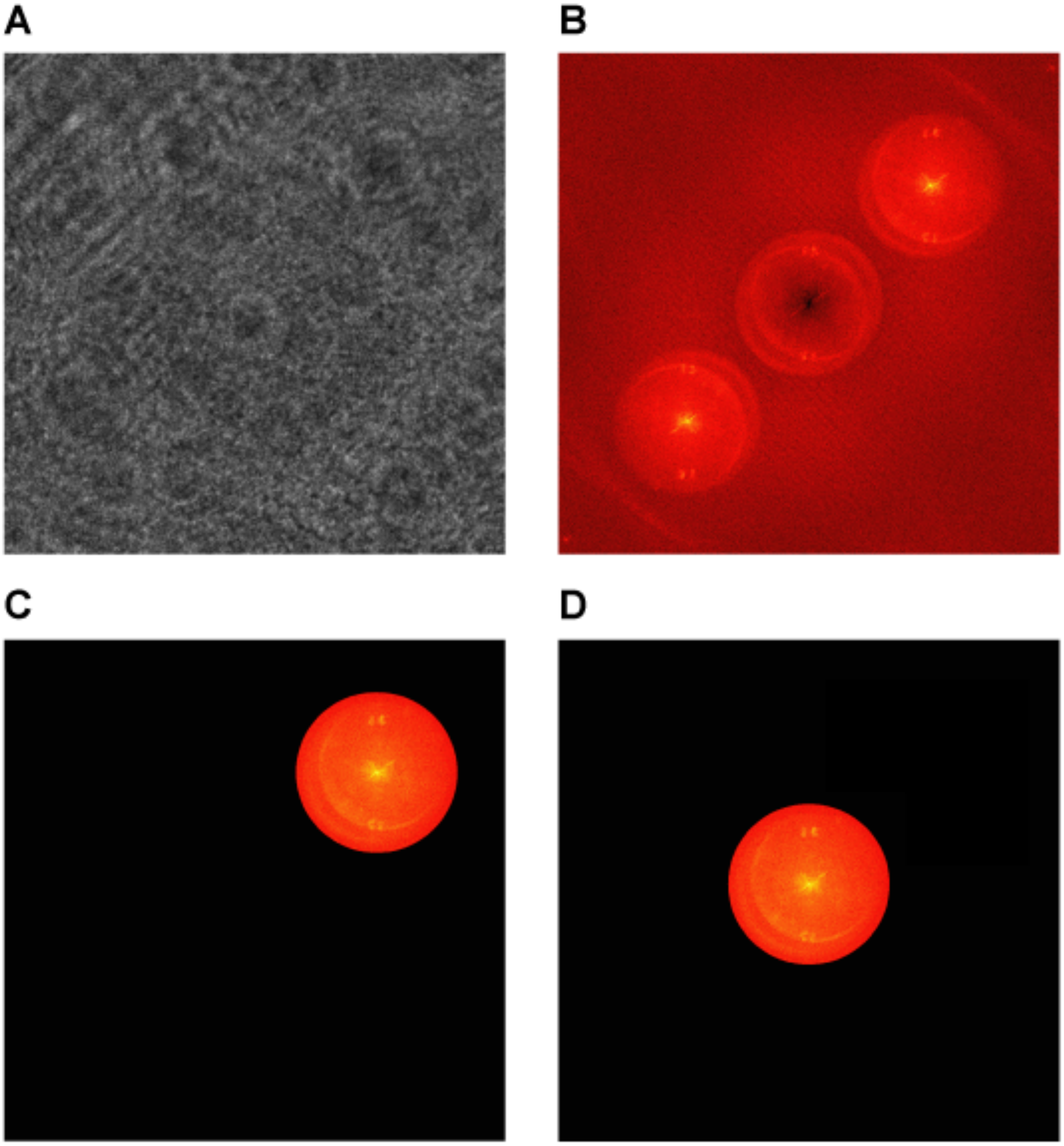


**Supplementary Figure 2 -** The workflow for the initial processing of holographic data. (a) A raw hologram. (b) The Fourier Transform of the raw hologram, plotted on a logarithmic scale. (c) The Fourier Transform of the hologram multiplied by a mask to isolate one of the high spatial frequency lobes. (d) The isolated lobe from (c) that has been linearly shifted to the center.

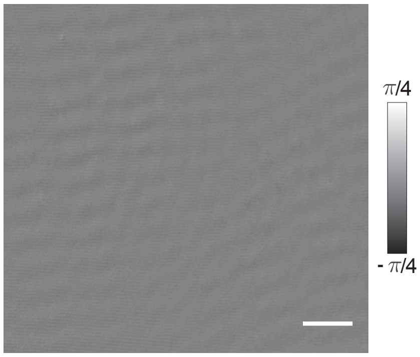


**Supplementary Figure 3 -** Representative DHM images of clustered GVs collapsed using 1.2 MPa of hydrostatic pressure. Scale bars represent 25 μm.
